# Identifying cardiac actinin interactomes reveals sarcomere crosstalk with RNA-binding proteins

**DOI:** 10.1101/2020.03.18.994004

**Authors:** Feria A. Ladha, Ketan Thakar, Anthony M. Pettinato, Nicholas Legere, Rachel Cohn, Robert Romano, Emily Meredith, Yu-Sheng Chen, J. Travis Hinson

**Affiliations:** University of Connecticut Health Center, Farmington, CT 06030, USA; The Jackson Laboratory for Genomic Medicine, Farmington, CT 06032, USA; Cardiology Center, UConn Health, Farmington, CT 06030, USA

## Abstract

**Rationale:** Actinins are actin cross-linkers that are ubiquitously expressed and harbor mutations in heritable diseases. The predominant cardiac actinin is encoded by *ACTN2*, which is a core structural component of the sarcomere Z-disk. ACTN2 is required for sarcomere function through complex interactions with proteins involved in sarcomere assembly, cell signaling and transcriptional regulation. However, there remain critical gaps in our knowledge of the complete and dynamic cardiac actinin interactome, which could reveal new insights into sarcomere biology.

**Objective:** We sought to examine the cardiac actinin interactome through sarcomere assembly in human cardiomyocytes.

**Methods and Results:** We utilized CRISPR/Cas9, induced pluripotent stem cell technology and BioID to reveal cardiac actinin protein interactions in human cardiomyocytes. We identified 324 cardiac actinin proximity partners, analyzed networks, and studied interactome changes associated with sarcomere assembly. We focused additional studies on unexpected actinin interactions with effectors with RNA-binding functions. Using RNA immunoprecipitation followed by sequencing, we determined that RNA-binding partners uncovered by actinin BioID were bound to gene transcripts with electron transport chain and mitochondrial biogenesis functions. Mammalian two-hybrid studies established that IGF2BP2, an RNA-binding protein associated with type 2 diabetes, directly interacted with the rod domain of actinin through its K Homology domain. IGF2BP2 was necessary for electron transport chain transcript localization to vicinal RNA-binding proteins, mitochondrial mass and oxidative metabolism. IGF2BP2 knockdown also impaired sarcomere function in a cardiac microtissue assay.

**Conclusions:** This study expands our functional knowledge of cardiac actinin, uncovers new sarcomere interaction partners including those regulated by sarcomere assembly, and reveals sarcomere crosstalk with RNA-binding proteins including IGF2BP2 that are important for metabolic functions.

## Introduction

Actinin proteins are ubiquitous spectrin family members that cross-link actin, and are important for myriad cell functions including adhesion, migration, contraction, and signaling^1^. The four human actinin isoforms have distinct expression profiles (non-muscle *ACTN1* and *ACTN4*; cardiac muscle *ACTN2*; and skeletal muscle *ACTN3*), but share homologous amino acid sequences that are organized into three structural domains—an actin-binding (AB) domain composed of two calponin-homology domains, a central rod domain containing four spectrin-like repeats (SR), and a calmodulin-like domain (CaM) containing two EF hand-like motifs^2^. While it is thought that actinin evolved initially to regulate the early eukaryotic actin-based cytoskeleton^3^, it has acquired more elaborate functions in vertebrates, including a mechanical role in the sarcomere, a specialized contractile system required for striated muscle function^3, 4^. Moreover, normal actinin function is important for multiple human organ systems as inheritance of *ACTN1, ACTN2*, and *ACTN4* mutations have also been linked to congenital macrothrombocytopenia, hypertrophic cardiomyopathy, and focal segmental glomerulosclerosis of the kidney, respectively^5-7^.

In the cardiac sarcomere, alpha actinin-2 (referred henceforth as actinin) is a major structural component of the Z-disk, where it is essential for sarcomere assembly and function^8^. Actinin regulates sarcomere assembly by providing a scaffold for protein-protein interactions (PPIs) such as with titin (TTN) and actin through CaM and AB domains, respectively^8-10^. Secondary to these interactions, the thin and thick filament sarcomere structures are organized to promote twitch contraction. It has also been observed that while the Z-disk is one of the stiffest sarcomere structures, actinin demonstrates remarkably dynamic and complex molecular interactions such as with the actin cytoskeleton that are necessary for normal myocyte function^11^. Moreover, actinin interaction partners have also been described to exit the sarcomere to execute critical roles in cell signaling, protein homeostasis and transcriptional regulation^12^. Secondary to the vast repertoire and dynamics of actinin molecular interactions, there remain extensive gaps in our knowledge of not only how actinin regulates sarcomere assembly and function, but also within other cellular contexts enriched for actinin expression such as at focal adhesions and cell surface receptors in both striated and non-striated cells that may inform how actinin mutations cause human disease^13, 14^.

Here, we examine the actinin interactome in CRISPR/Cas9-engineered human cardiomyocytes using proximity-dependent biotinylation (BioID). This study employs cardiomyocytes (iPSC-CMs) differentiated from induced pluripotent stem cells (iPSCs) that express actinin fused to the promiscuous biotinylating enzyme BirA* from the *ACTN2* locus to provide *in vivo* actinin expression levels and localization^15^. We identify a list of actinin neighborhood proteins including those that are dynamically regulated by sarcomere assembly to reveal molecular insights into this process. Our study uncovers unexpected actinin interactions with RNA-binding proteins including IGF2BP2 and specific RNA transcripts bound to these vicinal RNA-binding proteins including electron transport chain and mitochondrial biogenesis factors. We identify how this process is essential for electron transport chain and sarcomere contractile functions. Our study also provides a list of molecules that can be studied to reveal additional insights into sarcomere dynamics and unexpected functions of actinin.

## Methods

Detailed methods are provided in the Supplemental Materials. Statistical tests and samples sizes are described in each figure legend. The data that support the findings of this study are available from the corresponding author upon reasonable request.

### iPSC-CM Differentiation and CRISP/Cas9 studies

All stem cell experiments were performed using PGP1 iPSCs (Coriell #GM23338). iPSC-CMs were directly differentiated by sequential modulation of Wnt/β-catenin signaling, as previously described^16, 17^. For BioID studies, control and Actinin-BirA* iPSC-CMs were maintained in DMEM (Gibco 11965092) supplemented with homemade biotin-free B27 following metabolic enrichment^18, 19^. For proximity-labeling experiments, control and Actinin-BirA* iPSC-CMs were treated with 50 μM biotin for 24 hours followed by a 2-hour washout phase in biotin-free media before collection.

Genome editing was performed utilizing a CRISPR/Cas9 protocol adapted from previous studies^17, 20^. iPSCs were electroporated with pCas9-GFP (Addgene 44719), appropriate hU6-driven sgRNA, and HR targeting vector to generate *ACTN2-BirA\** knock-in iPSCs. Isogenic knockout of *TNNT2* was performed similarly, but in the absence of an HR vector and with GFP-based FACS enrichment in place of antibiotic selection.

### Streptavidin immunoprecipitation and Quantitative proteomics

After biotin washout, control and Actinin-BirA* iPSC-CMs were lysed and lysates were sonicated (Branson 250). Cleared lysates were added to magnetic Dynabeads MyOne Streptavidin C1 beads (Invitrogen 65001) and affinity purification was performed on a rotator overnight at 4°C. The following day, the beads were washed with appropriate buffers and washed beads were resuspended in PBS, snap frozen, and sent for quantitative proteomics to the Thermo Fisher Center for Multiplexed Proteomics (TCMP) at Harvard Medical School. For each tandem mass tag (TMT) mass spectrometry experiment, three biological replicates per condition were analyzed. For detailed methods regarding bead digestion, liquid chromatorgraphy-MS3 spectrometry (LC-MS/MS), and LC-MS3 data analysis please refer to full methods section.

TMT raw intensity values are provided for both experiments – experiment 1 (Supplemental Table 1) and experiment 2 (Supplemental Table 2). Before analysis, proteins previously reported to non-specifically bind BirA* were removed^21^. For analysis, TMT raw intensity values underwent Log2-fold-change (L2FC) calculation and enrichment analysis for Actinin-BirA* proteins relative to control. Significantly enriched proteins were those with L2FC ≥1 and false discovery rate (FDR) <0.05 (using two-way ANOVA followed by a two-stage linear step-up procedure of Benjamini, Krieger and Yekutieli to correct for multiple comparisons^22^). For visualization of data, Search Tool for the Retrieval of Interacting Genes/Proteins (STRING v. 11.0) was utilized for generation of interaction scores (proteins were not distinguished by isoform). Cytoscape (v.3.7.2) was utilized to generate interaction maps with an organic layout and the utilization of the CLUSTER and BINGO features for group clustering and appropriate GO term identification for each cluster. Quantitative proteomics data deposited to the ProteomeXchange Consortium identifier PXD018040.

### RNA-immunoprecipitation and sequencing (RIP-Seq)

After biotin washout, control and Actinin-BirA* iPSC-CMs were were lysed and cleared lysate was added to magnetic Dynabeads MyOne Streptavidin C1 beads (Invitrogen 65001). Samples and beads were incubated on a rotator overnight at 4°C. The following day, the beads were washed with appropriate buffer – all steps were performed in a 4°C room. Samples were then incubated with Proteinase K at 55°C for 30 minutes. Beads were resuspended in TRIzol (Invitrogen 15596018), RNA extraction was performed immediately followed by RNA-sequencing. RNA Samples were sent for sequencing at the University of Connecticut Institute for Systems Genomics. RNA sequencing libraries were generated using Illumina TruSeq Stranded Total RNA library preparation (Ribo-Zero depletion for rRNA was utilized for RIP-seq). Illumina NextSeq 500/550 sequencing was conducted with the v2.5 300 cycle reagent kit 9 (High Output). Estimated total single end reads per sample = 30-35M 150bp PE reads. Samples sequences were aligned with STAR to the hg38 human genome. In order to look at differential expression, DESeq2 (Bioconductor) was utilized. Genome sequencing datasets are deposited at GEO under accession numbers: GSE144806.

### Mammalian-Two Hybrid

Actinin (full-length and domains), IGF2BP2 (full-length and domains), PCBP1, PBCBP2, and SERBP1 were amplified from human cDNA and cloned into the appropriate vectors (pACT or pBIND) provided by the CheckMate Mammalian Two-Hybrid kit (Promega E2440). The appropriate combination of edited pACT and pBIND vectors, in addition to the kit-provided pGluc5 vector, were transfected using PEI into HEK 293T cells (ATCC CRL-3216), as outlined in the manufacturer’s protocol. In addition, the kit-provided positive and negative controls were transfected simultaneously for all experiments to verify proper signal production. Luciferase and Renilla activity were measured using manufacturer’s protocol (Promega E1910) utilizing BioTek’s Synergy 2 multi-mode microplate reader.

## Results

### BioID to identify actinin proximity partners

Actinin is an integral member of the actin cytoskeleton where it interacts within multiprotein complexes with diverse biophysical properties such as within Z-disks and membrane-associated focal adhesions, which are also dynamically regulated (Figure 1A). Secondary to these actinin properties, we selected the method of BioID that has been shown to be effective within these contexts^23^. BioID uses BirA*, which is a genetically-modified version of *E. coli* BirA that enables promiscuous lysine biotinylation within ∼10 nm radius of expression (Figure 1B)^24^.

**Figure 1.**
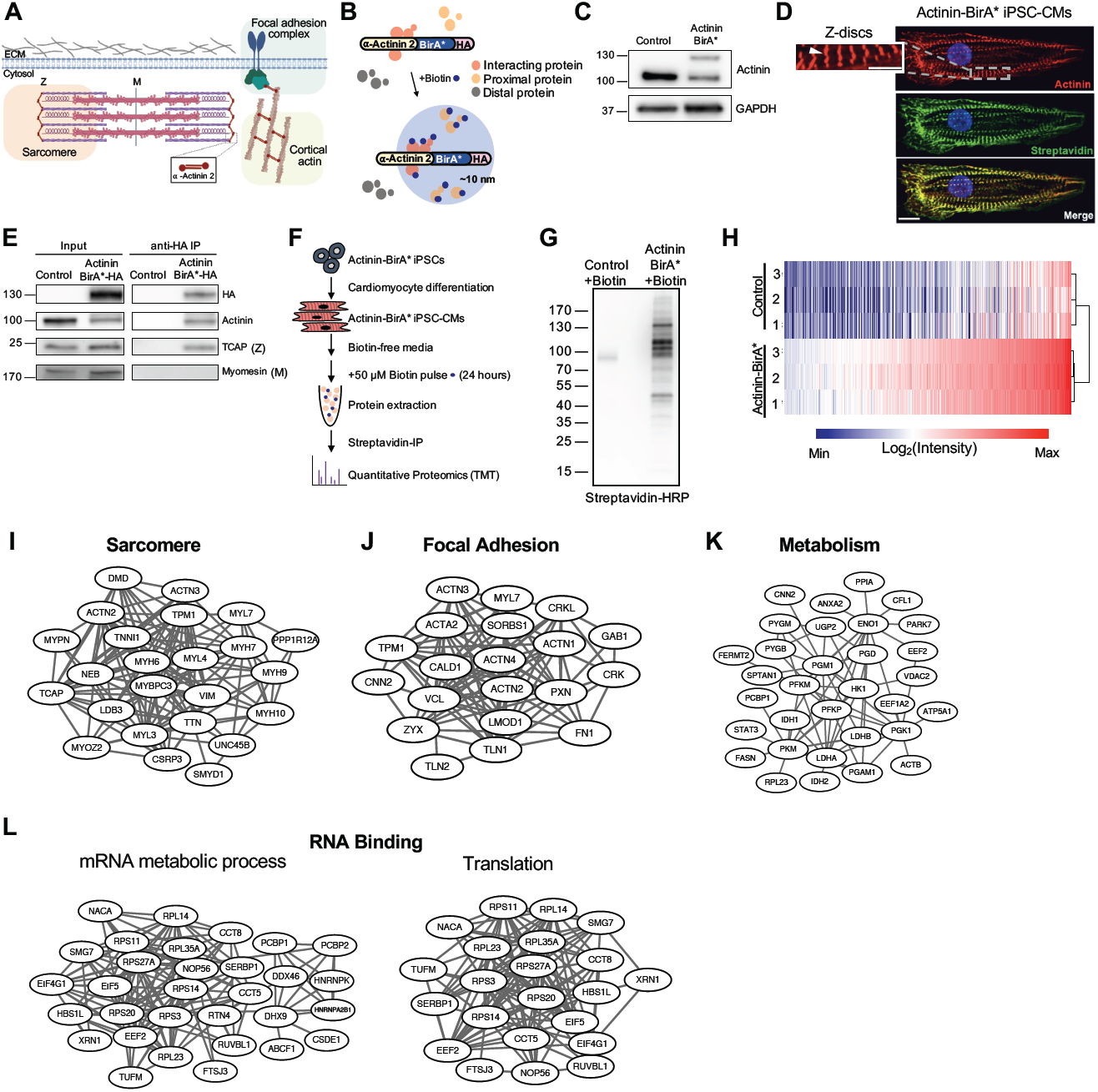
Actinin proximity interaction networks using BioID. **A)** Overview of actinin localization in cardiomyocytes including sarcomere Z-disk, focal adhesion and cortical actin cytoskeleton. **B)** Overview of the BioID method using Actinin-BirA* fusion to study actinin proximal protein networks. **C)** iPSC-CM lysates were probed with anti-actinin and anti-GAPDH to identify Actinin-BirA* fusion (∼130kD) and endogenous actinin (∼100kD). **D)** Confocal micrograph of Actinin-BirA* iPSC-CMs decorated with antibodies to actinin (red), streptavidin-AF488 (green) and DAPI DNA co-stain (blue) (scale bars, main image=10μm, inset=5μm). **E)** To validate localization, iPSC-CM lysates were immunoprecipitated with an anti-HA antibody, and probed with antibodies to known sarcomere components at Z-disk (anti-TCAP) and M-line (anti-myomesin), as well as anti-HA and anti-actinin controls. **F)** Overview of BioID experimental methods. **G)** iPSC-CM lysates were probed with streptavidin-HRP to examine Actinin-BirA*-biotinylated proteins. **H)** Heatmap and hierarchical clustering of Log2-transformed intensity values for the 324 significantly Actinin-BirA* enriched hits (L2FC ≥1 and FDR <0.05 relative to control) from combined TMT experiments. **I-L)** Interaction maps of TMT results based on STRING and GO terms – sarcomere, focal adhesion, metabolism, and RNA-binding.

We employed a genome engineering strategy to fuse BirA* with a hemagglutinin (HA) epitope tag to the carboxyl-terminus of endogenous *ACTN2* (Actinin-BirA*) in a human iPSC line using CRISPR/Cas9-facilitated homologous recombination (Supplemental Figure 1A). Following differentiation to iPSC-CMs, expression of the Actinin-BirA* fusion protein was confirmed (Figure 1C)^25, 26^. To verify that Actinin-BirA* functioned similarly to unmodified actinin, we performed several assays. We first utilized 3-dimensional cardiac microtissue assays that we previously employed to study sarcomere function by quantifying twitch force using cantilever displacement^17, 20^. Relative to unmodified actinin, twitch force was unaltered by Actinin-BirA* expression (Supplemental Figure 1B). We next confirmed appropriate localization of Actinin-BirA* to the Z-disk by both co-localization immunofluorescence (Figure 1D) and HA-immunoprecipitation assays (Figure 1E). Actinin-BirA* interacted with the Z-disk protein titin-cap (TCAP), but not the sarcomere M-line protein myomesin^27-30^, which additionally supported appropriate sarcomere localization. With the knowledge that Actinin-BirA* functions similarly to unmodified actinin, we next optimized biotin supplementation conditions in iPSC-CMs and confirmed that optimal labeling is achieved after 50 μM biotin supplementation (Figure 1F and Supplemental Figure 1C). Under these conditions, Actinin-BirA* biotinylated targets that overlapped actinin localization in iPSC-CMs (Figure 1D), and demonstrated abundant protein biotinylation compared to biotin-pulsed wildtype (WT) control iPSC-CMs (Figure 1G and Supplemental Figure 1D).

To identify and quantify actinin proximity proteins, we subjected both biotin-pulsed control and Actinin-BirA* iPSC-CM lysates to streptavidin immunoprecipitation and tandem mass tag (TMT) quantitative proteomics (Figure 1F). Analysis of Actinin-BirA* versus control samples identified enrichment of 324 actinin proximity partners (L2FC ≥1 and FDR <0.05; Supplemental Table 3), which are presented using hierarchical clustering and heatmap analysis (Figure 1H). The proximity list was next examined using Search Tool for the Retrieval of Interacting Genes/Proteins (STRING)^31^ network analysis (Supplemental Figure 1E). STRING confirmed identification of previously-known direct actinin interaction partners localized to the Z-disk including TTN^10^, TCAP^10^ and myozenin 2 (MYOZ2)^32^ (Figure 1I), which also belonged to a sarcomere cluster including numerous other sarcomere components determined by Gene-Ontology (GO) enrichment analysis. Actinin network proteins were also enriched within a focal adhesion cluster including previously-known direct actinin partner zyxin (ZYX)^33^ (Figure 1J). BioID also uncovered less well-described actinin proximity network clusters including metabolism (Figure 1K), and RNA-binding (Figure 1L).

### Actinin proximity partners change with sarcomere assembly

During sarcomere assembly, actinin first organizes into punctate Z-bodies, or pre-myofibrils, that coalesce to form striated Z-disks^34^. Z-disk assembly is critical to cardiomyocyte contractile function, but is incompletely understood. As it has been previously observed that cardiomyocytes that lack troponin T (cTnT; encoded by *TNNT2*) only assemble Z-bodies^35^, we studied cTnT knockout (KO) and wildtype (WT) iPSC-CMs as a model to understand actinin proximity partners through two stages of sarcomere assembly. We started by inserting a homozygous cTnT-KO mutation into the Actinin-BirA* iPSC line using CRISPR/Cas9 and utilized error-prone non-homologous end joining repair to generate an out-of-frame homozygous deletion mutation. We next verified that cTnT-KO iPSC-CMs expressed no cTnT protein (Supplemental Figure 2A), and actinin antibodies only decorated Z-body structures (Figure 2A). We then performed TMT to discern changes to actinin proximity partners between cTnT-WT and cTnT-KO iPSC-CMs (Supplemental Figure 2B). We identified 294 Actinin-BirA*-enriched proteins compared to control (L2FC ≥1 and FDR <0.05; Supplemental Table 4), which are presented using hierarchical clustering and heatmap analysis in Figure 2B. Using this subset of Actinin-BirA*-enriched proteins, we then analyzed cTnT-WT relative to cTnT-KO to assess enrichment with sarcomere assembly (Figure 2C). Overall, we identified 47 sarcomere assembly-enriched actinin proximity partners (L2FC ≥1 and FDR <0.05; Supplemental Table 4), which clustered into GO terms for “sarcomere” and “actin cytoskeleton” (Figure 2D).

**Figure 2.**
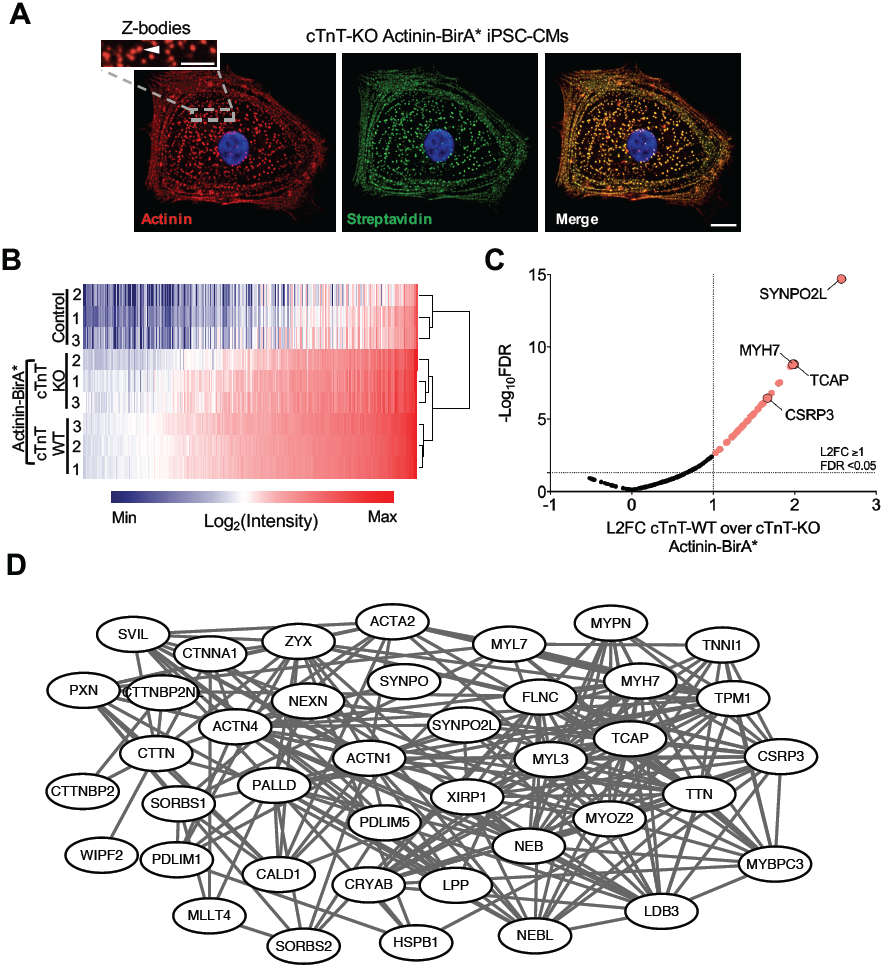
Sarcomere assembly-dependent actinin proximity interaction networks. **A)** Confocal micrograph of cTnT-KO Actinin-BirA* iPSC-CMs decorated with antibodies to actinin (red), streptavidin-AF488 (green), and DAPI DNA co-stain (blue) (scale bars, main image=10μm, inset=5μm). In contrast to the Z-disks observed in cTnT-WT iPSC-CMs (Figure 1D), cTnT-KO iPSC-CMs have punctate Z-bodies. **B)** Significant hits from TMT experiment 2 using hierarchical clustering and heatmap of Log2-transformed intensity values for the 294 Actinin-BirA*-enriched proteins compared to control (L2FC ≥1 and FDR <0.05). **C)** -Log_10_FDR (y-axis) plotted against the L2FC of cTnT-WT samples relative to cTnT-KO (x-axis) using the 294 Actinin-BirA*-enriched proteins, which identified 47 proteins (pink) that are further enriched with cTnT-dependent sarcomere assembly (L2FC ≥1 and FDR <0.05) **D)** STRING interaction map of sarcomere assembly-enriched TMT results analyzed by cluster analysis reveals sarcomere and actin cytoskeleton GO terms.

We next examined a prevailing hypothesis regarding sarcomere assembly that non-muscle myosin isoforms localize to Z-bodies, and are subsequently replaced by muscle myosin isoforms in Z-disks^34^. Our results confirm this model, as actinin proximity with cardiac muscle beta myosin (MYH7) is significantly enriched with Z-disk assembly in cTnT-WT iPSC-CMs (L2FC=1.976 and FDR <0.001), while non-muscle type B myosin (MYH10) is not (L2FC=0.055 and FDR=0.67) (Figure 2C). In addition, we also identified enrichment of late assembling proteins^36^ including telethon/titin-cap (TCAP; L2FC=1.993 and FDR<0.001). The most enriched sarcomere assembly partner was Synaptopodin 2-like (SYNPO2L; L2FC=2.573 and FDR<0.001), which is a known binding partner of actinin that is critical to heart and skeletal muscle development, as disruption in zebrafish results in disorganized sarcomere structure and impaired contractility^37^. Another actinin partner of note was cysteine and glycine rich protein 3 (CSRP3; L2FC=1.666 and FDR<0.001). CSRP3 is a Z-disk protein that has been implicated to regulate mechanical stretch signaling^38^, anchor calcineurin to the sarcomere^39^, and cause heart failure when mutated^40^. In summary, we utilized Actinin-BirA* proximity proteomics in a two-stage sarcomere assembly model, which identified enrichment of specific interactions at late Z-disks relative to early Z-bodies, including differentially-regulated muscle and non-muscle myosins, late assembling proteins, and additional Z-disk partners implicated in heart failure.

### Identification of transcripts bound to actinin-proximal RNA-binding proteins

To reveal functional insights into actinin interactions with RNA-binding proteins, we next sought to identify gene transcripts bound to RNA-binding proteins biotinylated by Actinin-BirA*. To do this, we performed RNA immunoprecipitation followed by sequencing (RIP-seq) (Figure 3A)^41^. Principle component analysis (PCA) demonstrated clustering of Actinin-BirA* transcripts compared to controls (Figure 3B), and differential expression analysis identified 945 transcripts bound to biotinylated RNA-binding proteins (Supplemental Table 5). GO analysis (Figure 3C) revealed enrichment of electron transport chain (ETC) and ribosome transcripts (Figure 3D and Supplemental Figure 3A). In contrast, we observed no enrichment for glycolysis transcripts (Figure 3E and Supplemental Figure 3B). 1113 transcripts with functions in chromatin and nuclear functions were depleted (Figure 3C). Cytoskeletal transcripts, including those encoding thin and thick filament sarcomere proteins, were also depleted (Figure 3F and Supplemental Figure 3C). In summary, RIP-seq demonstrated that RNA-binding proteins that were in proximity to cardiac actinin bind specific ETC and ribosome transcripts.

**Figure 3.**
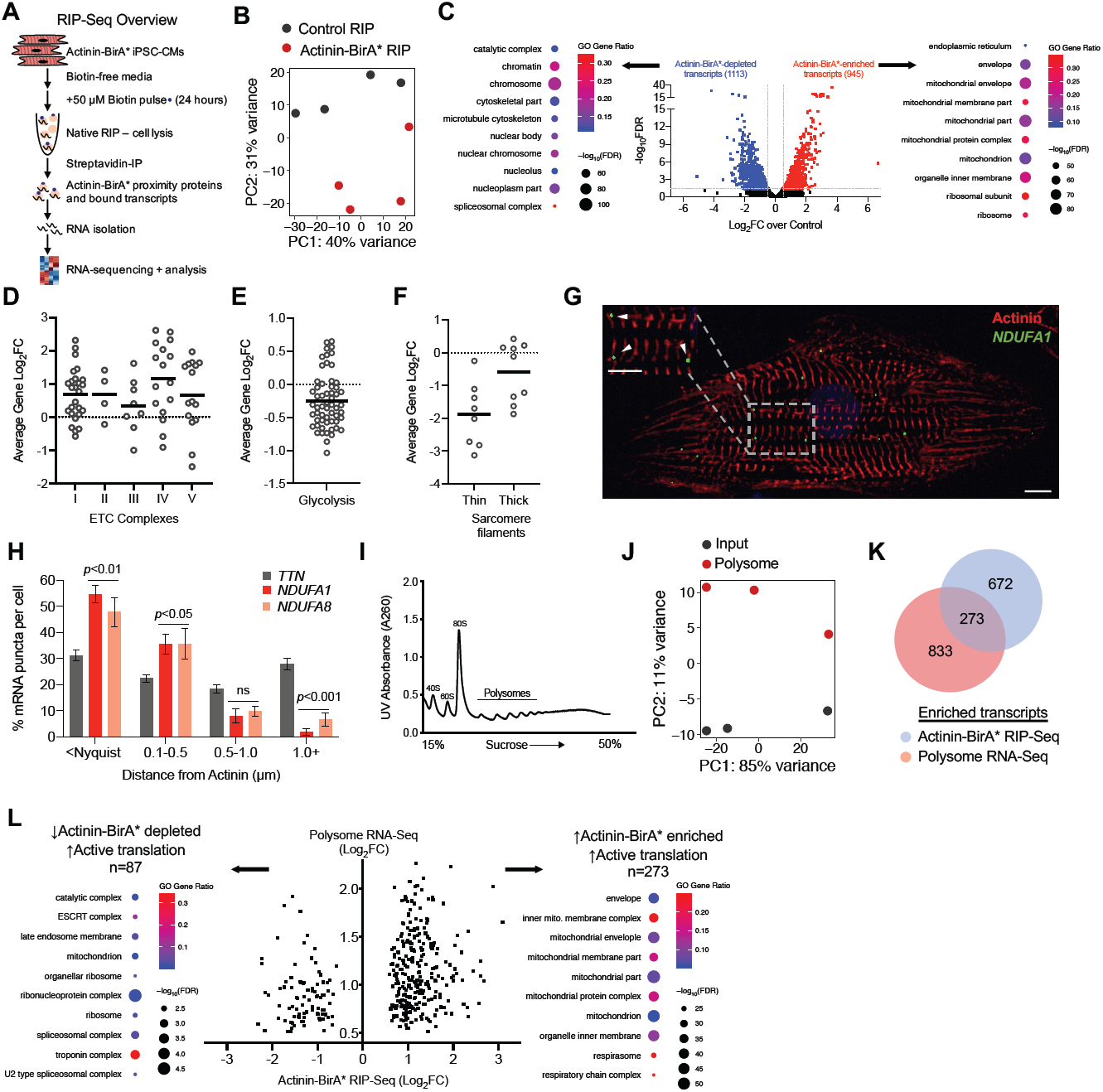
Identifying gene transcripts bound to actinin-proximal RNA-binding proteins using RIP-seq. **A)** Overview of modified RIP-seq protocol to study RNA transcripts bound to actinin-proximal RNA-binding proteins by streptavidin affinity purification of RNA-binding proteins followed by RNA isolation and sequencing. **B)** PCA plot shows clustering of Actinin-BirA* RIP-seq samples relative to controls (n=4). **C)** Volcano plot of RIP-seq data shaded red for 945 enriched transcripts (FDR <0.05, L2FC ≥0.5) and blue for 1113 depleted transcripts (FDR <0.05, L2FC ≤-0.5). Differentially-enriched transcripts were analyzed by GO and enrichment terms are listed for enriched or depleted samples. **D)** ETC gene components spanning respiratory chain complex I-V proteins are enriched on average. Each unshaded circle represents a single ETC gene. **E)** Glycolysis gene components are not enriched on average. Each unshaded circle represents a single glycolysis gene. **F)** Sarcomere gene components divided by localization to the thick or thin filament are not enriched or depleted on average. Each unshaded circle represents a single sarcomere gene. **G)** Representative confocal micrograph of iPSC-CMs decorated with antibodies to actinin (red), DAPI DNA co-stain (blue), and RNA FISH probes against *NDUFA1* (green). (scale bars, main image=10μm, inset=5μm). **H)** Quantification of RNA proximity to actinin protein. Data are derived from n=10 cells. **I)** Representative polysome profile of iPSC-CMs used for Polysome-seq studies. **J)** PCA plot shows clustering of polysome-enriched transcripts compared to controls. **K)** Venn diagram illustrating unique and overlapping transcripts between Actinin-BirA* RIP-seq and Polysome-seq (FDR <0.05, −0.5 ≤ L2FC ≥ 0.5 for both). **L)** Overlay of Actinin-BirA* RIP-seq and Polysome-seq results to identify highly translated genes (FDR <0.05, L2FC ≥0.5 by Polysome-seq) that are RIP-seq enriched (FDR <0.05, L2FC ≥0.5 by RIP-seq), or RIP-seq depleted (FDR <0.05, L2FC ≤-0.5 by RIP-seq) and the resulting GO terms for each gene set. Data are mean ± SEM; significance assessed by one-way ANOVA using Holm-Sidak correction for multiple comparisons (H).

Following qPCR validation of ETC transcript enrichment identified by RIP-seq (Supplemental Figure 3D), we performed RNA fluorescence *in situ* hybridization (RNA FISH) and Imaris distance quantification (Supplemental Figure 3E) to test transcript proximity relative to actinin protein expression. We found that RNA puncta encoding ETC component *NDUFA1* were more likely to be localized near actinin protein compared to the sarcomere thick filament transcript *TTN* (Figure 3G, 3H, and Supplemental Figure 3F). We also validated an additional actinin proximity protein-enriched transcript *NDUFA8* (Figure 3H, Supplemental Figure 3G). We next assessed co-localization between *NDUFA1* and *NDUFA8* transcripts, which could reflect potential ETC transcript storage or processing mechanisms. While both *NDUFA1* and *NDUFA8* transcripts localized near actinin protein, they did not overlap with each other (Supplemental Figure 3H).

With next performed polysome profiling (Figure 3I) followed by sequencing (Polysome-seq)^42^ to assess for the relationship between transcript translation and RIP-seq enrichment. PCA analysis demonstrated clustering of replicates of polysome-bound transcripts compared to controls (Figure 3J), and 273 transcripts were shared between the polysome-enriched and RIP-seq-enriched fractions (Figure 3K and Supplemental Table 6), suggesting that only a subset of RIP-seq-enriched transcripts may be actively translated. Shared transcripts represented 28.9% of RIP-seq-enriched transcripts, and 24.7% of total polysome-enriched transcripts. GO term analysis of overlapped transcripts revealed mitochondrial functions, which included ETC members (Figure 3L). Conversely, RIP-seq-depleted but polysome-enriched transcripts enriched for distinct GO terms including ribosome and troponin complex (Figure 3L). Compared to the overall iPSC-CM transcriptome, these data support that ETC transcripts, which were identified by RIP-seq as bound by RNA-binding proteins in proximity to actinin, are also more likely to be translated into protein.

### Actinin rod domains directly interact with IGF2BP2 K homology domains

Direct actinin interactions with RNA-binding proteins have not been previously well-described. As ETC transcripts function in oxidative phosphorylation, and were RIP-seq enriched, we additionally studied candidate actinin-interacting RNA-binding proteins secondary to their association with metabolic functions, including IGF2BP2^43^, PCBP1^44^, PCBP2^45^, and SERBP1^46^. We first confirmed actinin proximity results using streptavidin affinity purification followed by immunoblotting against the four RNA-binding proteins (Figure 4A and Supplemental Figure 4A). To validate that actinin and candidate RNA-binding proteins interact, we also confirmed co-immunoprecipitation of Actinin-BirA* using an HA epitope antibody with IGF2BP2, PCBP1 and PCBP2 (Figure 4B and Supplemental Figure 4B). The lack of SERBP1 co-immunoprecipitation suggested a more complex actinin interaction mechanism or a false positive result.

**Figure 4.**
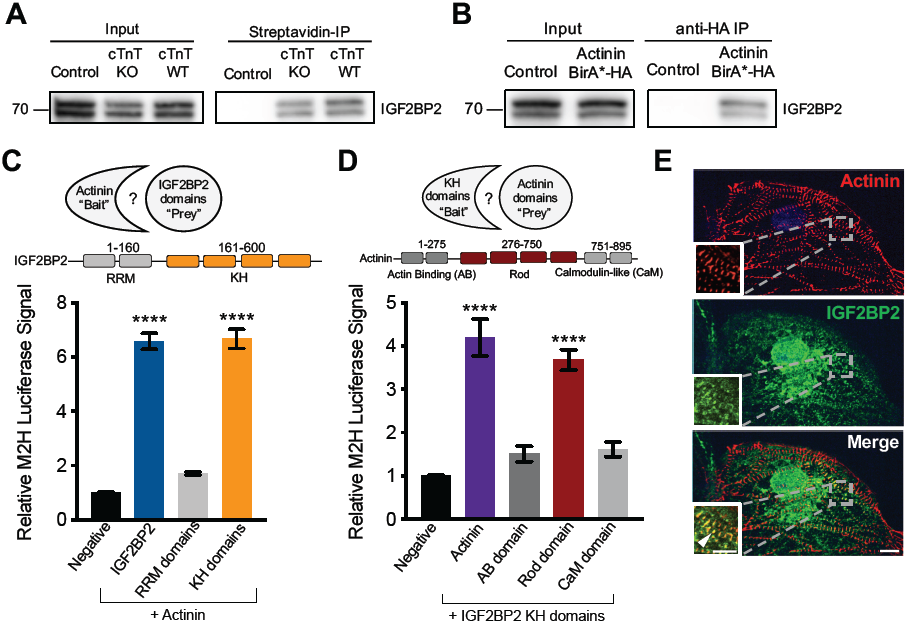
RNA-binding protein IGF2BP2 directly binds actinin. **A)** Representative immunoblot probed with antibodies to IGF2BP2 in streptavidin affinity-purified lysates. Double band represents two IGF2BP2 splice products. **B)** Representative immunoblot probed with antibodies to IGF2BP2 in anti-HA immunoprecipitated lysates. Double band represents two IGF2BP2 splice products. **C)** Mammalian-two-hybrid (M2H) study conducted in HEK 293T cells to fine map the direct interaction between actinin and IGF2BP2. IGF2BP2 was divided into RRM (residues 1-160) and KH (residues 161-600) domains to fine-map the interaction. All data are n=3. **D)** M2H results to fine map the direct interaction between the KH domain of IGF2BP2 and Actinin. Actinin was divided into AB (residues 1-275), rod (residues 276-750) and CaM domains (residues 751-895). All data are n=3. **E)** Representative confocal micrograph demonstrating partial co-localization of actinin (anti-actinin; red) with IGF2BP2 (anti-IGF2BP2; green) with DAPI DNA (blue) co-stain in iPSC-CMs. (scale bars, main image=10μm, inset=5μm). Data are mean ± SEM; significance assessed by one-way ANOVA using Holm-Sidak correction for multiple comparisons (C, D) and defined by *P* ≤ 0.0001 (****).

To extend co-immunoprecipitation results, we also tested the mode of interaction between actinin and IGF2BP2, PCBP1, PCPB2, and SERBP1 using mammalian 2-hybrid (M2H) assays in HEK 293T cells. We began by constructing expression vectors encoding full-length or structural domains of actinin and RNA-binding proteins. M2H using IGF2BP2 as “prey” and full-length actinin as “bait” resulted in luciferase activation, but not for PCBP1, PCBP2 or SERBP1 (Figure 4C and Supplemental Figure 4C). We next mapped the IGF2BP2-actinin interaction to IGF2BP2’s K Homology (KH) structural domain but not its RNA recognition motif (RRM). Using IGF2BP2 as “bait”, we then mapped the actinin-binding site to its rod domain (Figure 4D). In support of these findings, we observed co-localization of IGF2BP2 with actinin by immunofluorescence at Z-disk subsets (Figure 4E). Taken together, these data (Supplemental Figure 4D) both validate our BioID results that implicate actinin as an interaction hub for RNA-binding proteins, and establish that IGF2BP2 directly binds actinin through specific structural domains.

### IGF2BP2 regulates ETC transcript localization, oxidative phosphorylation, and contractile function in cardiac microtissues

IGF2BP2 has been shown to bind ETC transcripts in glioblastoma stem cell lines^47^, and this observation provided the rationale to test whether IGF2BP2 could also be responsible for the RIP-seq enrichment of ETC transcripts that we observed in Actinin-BirA* samples. We employed a genetic knockdown approach using an shRNA that decreased IGF2BP2 protein levels in iPSC-CMs (Figure 5A), and performed RIP-qPCR in IGF2BP2-depleted iPSC-CMs focused on ETC transcripts *NDUFA1, NDUFA8, NDUFA11*, and *COX6B1*. IGF2BP2 knockdown decreased all affinity purified ETC transcripts (Figure 5B) but not total cellular transcript levels (Figure 5C)—suggesting that IGF2BP2 is necessary for RIP-seq enrichment but not overall transcript stability. To validate results by a second method, we also sequentially precipitated transcripts by IGF2BP2 antibody first to collect all IGF2BP2-bound transcripts, followed by streptavidin affinity to study the subset of IGF2BP2-bound transcripts that have been biotinylated secondary to actinin proximity (Supplemental Figure 5A). We confirmed ETC transcript enrichment at biotinylated IGF2BP2 (Supplemental Figure 5A), and concluded that RIP-seq enrichment of ETC transcripts is dependent on IGF2BP2 levels.

**Figure 5.**
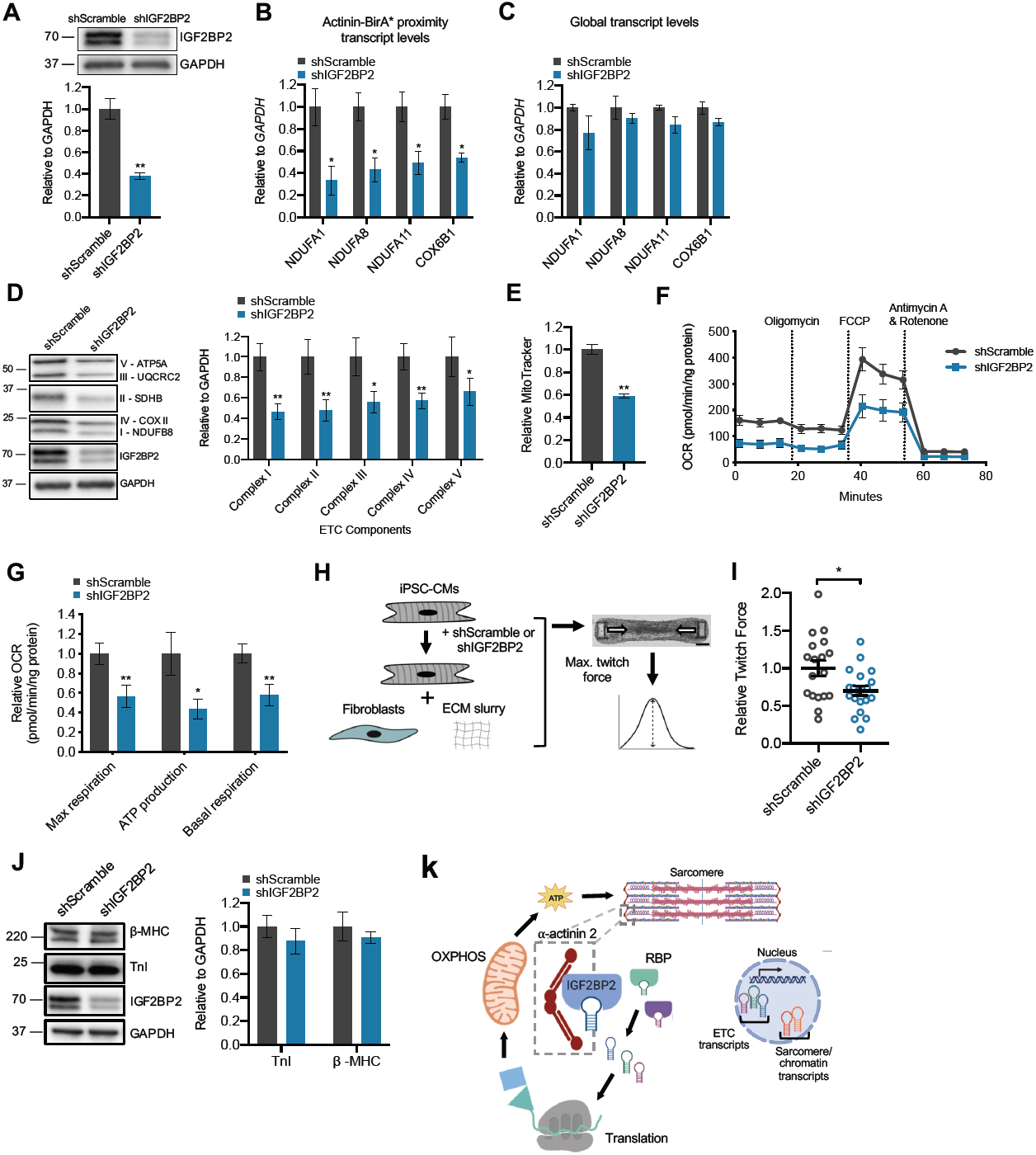
IGF2BP2 regulates ETC transcript localization, oxidative phosphorylation, and contractile function in cardiac microtissues. **A)** Representative immunoblot of lysates from iPSC-CMs treated with shRNA targeting IGF2BP2 or scramble control probed with antibodies to IGF2BP2 and GAPDH. Double band represents two IGF2BP2 splice products. **B)** Streptavidin affinity purification of transcripts bound to actinin proximal RNA-binding proteins after IGF2BP2 knockdown, followed by qPCR of ETC transcripts *NDUFA1, NDUFA8, NDUFA11* and *COX6B1*. **C)** QPCR analysis of global transcript levels of *NDUFA1, NDUFA8, NDUFA11* and *COX6B1* after IGF2BP2 knockdown. **D)** Representative immunoblot of lysates from iPSC-CMs after IGF2BP2 knockdown, and probed with antibodies to ETC complex I-V subunits (anti-ATP5A, UQCRC2, SDHB, COX II and NDUFB8), IGF2BP2 and GAPDH control. **E)** FACs analysis of iPSC-CMs stained with MitoTracker after IGF2BP2 knockdown (N>5000 events per replicate). **F)** Oxygen consumption rates normalized to protein levels after IGF2BP2 knockdown in iPSC-CMs, and **(G)** specific OCR parameter quantification. **H)** Overview of cardiac microtissue platform and twitch force analysis. ECM-extracellular matrix slurry. Fibroblasts-normal human ventricular fibroblasts. (scale bar-150μm). **I)** Cardiac microtissues exhibit reduced twitch force after IGF2BP2 knockdown. Data are n≥10 microtissues. **J)** Representative immunoblot of lysates from iPSC-CMs probed with antibodies to myosin heavy chain 7 (β-MHC) and troponin I (TnI) proteins show no change in expression after IGF2BP2 knockdown. Anti-GAPDH used as additional control. **K)** Proposed model of an actinin-based PPI hub that regulates oxidative phosphorylation (OXPHOS) through proximity interactions with RNA binding proteins (RBP) including IGF2BP2 and translational machinery. Data are n≥3; mean ± SEM; significance assessed by Student’s t-test (A-E, G, I, J) and defined by *P* <0.05 (*), *P* ≤ 0.01 (**).

We next studied the IGF2BP2-dependence of ETC protein quantity and oxidative phosphorylation. To do this, we performed IGF2BP2 knockdown followed by protein level quantification of subunits from ETC complexes I-V as well as the glycolytic control, GAPDH. IGF2BP2 knockdown diminished protein levels of components of all five ETC complexes, but not GAPDH (Figure 5D). Because MTCO2, a complex IV subunit that is transcribed and translated in mitochondria^48^, this result suggested that IGF2BP2 knockdown may have also reduced mitochondrial mass in iPSC-CMs. In support of this, we found reduced MitoTracker Green FM signal (Figure 5E) and oxygen consumption rates in IGF2BP2 knockdown iPSC-CMs relative to controls (Figure 5F, 5G). Re-analysis of our RIP-seq data also identified enrichment for *ESRRA* (Supplemental Table 6A), encoding estrogen-related receptor alpha that is an essential regulator of cardiac mitochondrial biogenesis, which could contribute to the reduction in mitochondrial mass^49^.

As cardiomyocytes rely on oxidative metabolism^50^, we next hypothesized that IGF2BP2 knockdown could impair energy-consuming functions such as sarcomere contraction. To test sarcomere function, we performed IGF2BP2 knockdown in iPSC-CMs and produced cardiac microtissues (Figure 5H). Compared to controls, IGF2BP2 knockdown resulted in decreased twitch force (Figure 5I). We confirmed that diminished twitch force was not secondary to changes in sarcomere protein levels as beta myosin heavy chain (β-MHC) and troponin I (TnI) were unchanged (Figure 5J). In summary, we propose the model that actinin functions as a PPI hub that connects the sarcomere Z-disk to electron transport chain regulation through direct interactions with IGF2BP2 and other RNA-binding partners that are essential to support normal cardiomyocyte contractile function (Figure 5K).

## Discussion

The actinins are multipurpose actin cross-linkers that execute their tasks through PPIs with myriad partners with diverse biophysical properties and kinetics including components of the cardiac sarcomere. Here, we employed quantitative proteomics and BioID to understand the identities, organization and functions of cardiac actinin PPI hubs using proximity-dependent biotinylation in human iPSC-CMs. The number of actinin interactions identified in our study was made possible by the unique capabilities of BioID including the ability to detect both direct and vicinal interactions, static and dynamic interactions and interactions in diverse microenvironments such as with membrane-associated partners. In contrast, previous actinin PPI studies have generally utilized yeast two-hybrid approaches that can be challenging with proteins that dimerize and can self-activate yeast reporters^51^. The robustness of our interaction study was achievable through fusion of BirA* into the endogenous actinin locus made possible with CRISPR/Cas9, which led to Actinin-BirA* expression levels and localization that mimicked unaltered actinin. By targeting the endogenous genetic locus, our approach could also be extended to studies of particularly large proteins including other spectrin family proteins, or to other cell lineages differentiated from iPSCs to understand cell type-specific PPIs.

A major objective of this study was to determine the molecular composition of the Z-disk through sarcomere assembly. Our current understanding of Z-disk proteins is predominantly derived from stitching together smaller PPI studies focused usually on a single interaction, but has not been systematically characterized, especially through a dynamic process such as sarcomere assembly. Here, we combined BioID with a two-stage sarcomere assembly model (i.e. Z-body and Z-disk) and confirmed previous Z-disk proteins including TTN^10^, TCAP^32^, LDB3^52^, MYOZ2^32^, CSRP3^39^, and SYNPO2L^37^, demonstrating the specify of our approach. This enabled us to capture enrichment of muscle myosin isoforms and late assembling proteins, which confirmed and clarified principles of previous myofibrillogenesis models^34^. Future studies may also be enabled by our actinin interactome study to inform new myofibrillogenesis mechanisms and undescribed consequences of disease-associated mutations within Z-disk molecules such as *ACTN2* itself.

Most importantly, our study yielded unexpected actinin functions in the regulation of RNA biology through interactions with RNA-binding proteins that we demonstrate to be important for the control of oxidative metabolism and mitochondrial mass. In zebrafish, 71% of transcripts are localized in spatially-distinct patterns^53^, which provides spatial and temporal regulation of gene expression such that local stimuli can regulate translation on-site^54^. The sarcomere itself has not been well-understood to regulate RNA localization and functions, but we find that actinin serves as a PPI hub for RNA-binding proteins and translational machinery, which we exploited to uncover a spatial relationship between ETC transcripts and actinin expression. Our study provides potential new mechanisms for the previous observation that intermyofibrillar mitochondria both reside adjacent to the Z-disk^55^, and have higher rates of oxidative phosphorylation^56^. We observe that ETC transcripts are in proximity to actinin protein that is dependent on IGF2BP2, an RNA-binding protein associated with type 2 diabetes by genome-wide association studies^43^. In our study, IGF2BP2 knockdown using a 3-dimensional cardiac microtissue model resulted in diminished contractile function in association with reduced ETC protein levels, mitochondrial mass and oxygen consumption rates in iPSC-CMs. Furthermore, this mechanism may inform why IGF2BP2 knock-out mice also have impaired fatty acid oxidation in skeletal muscle where actinin is also highly expressed^57^ or new insights into mechanisms of diabetic cardiomyopathy.

Our study has important limitations including the use of human iPSC-CMs. These cells empowered the opportunity to study sarcomere assembly in a human cellular context, but do not achieve the maturity of adult cardiomyocytes. In the future, engineering and investigating mouse models with Actinin-BirA* could further refine and clarify Z-disk interactions in the healthy heart and in diseases such as heart failure. Our study could also be extended to employ more dynamic approaches such as with the recently-described TurboID enzyme^58^ which exhibits improved biotinylation kinetics and labeling efficiencies compared to BirA*. Our RIP-seq approach is also limited by the inability to directly label actinin-proximal transcripts, which could be enabled by the recently-developed method of APEX-RIP^49^. While we tested IGF2BP2 functions by genetic knock-down that is limited by the inability to exclude actinin-independent IGF2BP2 functions, we were not able to fine-map the subdomain interaction that could be altered to more precisely establish the functional relevance of this single interaction. This is consistent with previous observations that spectrin-family rod domain interactions require higher-order tertiary and quaternary structures for PPI^1^.

In summary, we provide a catalogue of actinin proximity interactions including those involved in sarcomere assembly such as new molecules not previously implicated in sarcomere biology such as RNA-binding factors. We identify how interactions between cardiac actinin and RNA-binding proteins including IGF2BP2 regulate cardiomyocyte oxidative phosphorylation, which is a new mode of metabolic regulation spatially localized to the energy-consuming sarcomere. Further exploration and refinement of these interactions could reveal new therapeutic targets and pathophysiology for diseases of the cardiomyocyte including heart failure.

## Supporting information

Control tissue movie

## Acknowledgements

This study could not have been completed without the assistance of Bo Reese (UConn Center for Genome Innovation) for RNA sequencing and RIP-seq, and Anthony Carcio (The Jackson Laboratory for Genomic Medicine) for flow cytometry. TMT proteomics and quantitative analyses was supported by Sanjukta Thakurta from the Thermo Fisher Center for Multiplexed Proteomics at Harvard Medical School. Artwork was created using BioRender. Author contributions: Investigation and Validation, F.A.L., A.M.P., K.T., N.L., and J.T.H.; Cells and/or reagent generation, F.A.L., A.M.P., K.T., N.L., R.C., R.R., E.M., Y.S.C., and J.T.H.; Formal Analysis, F.A.L., A.M.P., and J.T.H.; Funding Acquisition, F.A.L., A.M.P., and J.T.H.; Conceptualization and Supervision, J.T.H; Writing, F.A.L., A.M.P., and J.T.H. All authors reviewed the manuscript prior to submission.

## Sources of Funding

Funding for this study was obtained from the National Institutes of Health (J.T.H., HL125807, HL142787, and EB028898), American Heart Association (F.A.L., PRE35110005, and A.M.P., PRE34381021) and institutional start-up funds (UConn Health).

## Disclosures

All authors declare no conflicts of interest regarding the work presented here.

## Notes

### Competing Interest Statement

The authors have declared no competing interest.

