## Supplementary Material for "Identifying cardiac actinin interactomes reveals sarcomere crosstalk with RNA-binding proteins"

#### Expanded Methods

##### iPSC Culture and iPSC-CM Differentiation

All stem cell experiments were done in PGP1 iPSCs (Coriell #GM23338), which have been extensively characterized including whole genome sequencing and karyotyping as part of the ENCODE project (ENCBS368AAA). iPSCs were maintained on Matrigel-coated tissue culture plates (Corning 354230) in mTeSR1 (STEMCELL Technologies 85875). iPSCs were passaged at 80-90% confluency utilizing Accutase (BD 561527) and 10 μM ROCK inhibitor Y-27632 (Tocris 1254). iPSC-CMs were directly differentiated by sequential modulation of Wnt/β-catenin signaling, as previously described[1, 2]. On Day 13 of differentiation, iPSC-CMs were purified by metabolic enrichment using glucose-free DMEM (Gibco 11966025) supplemented with 4 mM lactate (Sigma 71718) for 24-48 hours[3]. Following selection, iPSC-CMs were trypsinized (Gibco 25200056) and re-plated onto fibronectin-coated tissue culture plates (Gibco 33016015). iPSC-CMs were maintained in RPMI-B27. For proximity-labeling experiments, control and Actinin-BirA* iPSC-CMs were maintained in DMEM (Gibco 11965092) supplemented with homemade biotin-free B27 following metabolic enrichment[4, 5]. For BioID studies, control and Actinin-BirA* iPSC-CMs were treated with 50 μM biotin for 24 hours followed by a 2-hour washout phase in biotin-free media before collection. All iPSC-CM analyses were performed on Day 25-30 unless noted otherwise.

##### Plasmid Cloning

The homologous recombination (HR) targeting vector (System Biosciences HR220PA-1) was modified prior to being used in genome-editing experiments. A gene fragment for

BirA* (R118G)-HA tag was obtained (IDT) and cloned into the parent vector via Gibson assembly (NEB E2621). *ACTN2* ~800bp 5' and 3' HR arm gene fragments (IDT) were then cloned into the appropriate cloning sites (sequences in Table S5). All HR vector propagation steps were performed in DH5α *E.coli* (NEB C2987). For hairpin (shRNA) experiments, the pLKO.1 puro lentiviral plasmids containing either shScramble (Addgene 1864) or shIGF2BP2 (sequence in Table S5) were assembled using T4 DNA ligase (NEB M0202S) and propagated in NEB Stable *E.coli* (NEB C3040).

**CRISPR/Cas9 studies**

Genome editing was performed utilizing a CRISPR/Cas9 protocol adapted from previous studies[2, 6]. $8x10^6$ iPSCs were electroporated with 20 µg pCas9-GFP (Addgene 44719), 20 µg of the appropriate hU6-driven sgRNA (designed using crispr.mit.edu), and 20 µg of the HR targeting vector to generate *ACTN2-BirA** knock-in iPSCs. Electroporated cells were transferred to a Matrigel-coated 100 mm dish containing mTeSR1 and 10 µM Y-27632. The following day, selection was started with 50 µg/mL Hygromycin B (Invitrogen 10687010) to isolate single iPSC clones, which were then manually picked, expanded, and screened via Sanger sequencing. Isogenic knockout of *TNNT2* was performed similarly, but in the absence of an HR vector and with GFP-based FACS enrichment in place of antibiotic selection.

**Immunoprecipitation Experiments**

Streptavidin immunoprecipitation

After biotin washout, control and Actinin-BirA* iPSC-CMs were rinsed three times with room temperature PBS. Cells were lysed with lysis buffer (50 mM Tris-Cl pH7.4, 500 mM NaCl, 0.2% SDS, 1x protease inhibitor, 1mM DTT, $ddH_2O$). Lysates were sonicated

(Branson 250) two times on ice with 2 minutes between each cycle (30 pulses, 30% duty cycle, output level3). Samples were centrifuged to pellet cell debris and cleared lysate were added to magnetic Dynabeads MyOne Streptavidin C1 beads (Invitrogen 65001), which were prewashed with PBS and lysis buffer. Immunoprecipitation was performed on a rotator overnight at 4°C. The following day, the beads were resuspended in wash buffer 1 (2% SDS, $ddH_2O$), rotated for 8 minutes at room temperature, and then re-collected using a magnetic separation stand. The same steps were then repeated with wash buffer 2 (0.1% deoxycholic acid, 1% triton X-100, 1mM EDTA, 500 mM NaCl, 50 mM HEPES pH7.5, $ddH_2O$) and wash buffer 3 (0.5% deoxycholic acid, 0.5% NP-40, 1 mM EDTA, 250 mM LiCl, 10 mM Tris-Cl pH7.4, $ddH_2O$). Finally, the washed beads were resuspended in PBS, snap frozen, and sent for quantitative proteomics.

HA immunoprecipitation

Control and Actinin-BirA* iPSC-CMs were collected at Day 25 and HA-tag IP was conducted using reagents supplied in HA-tag IP/Co-IP Application set (Thermo Scientific 26180Y). The subsequent protein immunoblots were conducted as outlined in the immunoblot section.

RNA Immunoprecipitation (RIP)

After biotin washout, control and Actinin-BirA* iPSC-CMs were rinsed three times with room temperature PBS. Cells were lysed with ice-cold RIP buffer (150 mM KCl, 25 mM Tris-Cl pH7.4, 5 mM EDTA, 0.5 mM DTT, 0.5% NP40, 1x protease inhibitor, 100 U/ml RNase inhibitor, RNase-free water). Cell debris was pelleted and cleared lysate was added to magnetic Dynabeads MyOne Streptavidin C1 beads (Invitrogen 65001), which were prewashed with PBS and RIP buffer. Samples and beads were incubated on a rotator overnight at 4°C. The following day, the beads were washed three times with ice-cold RIP buffer – all steps were performed in a 4°C room. Samples were then incubated with Proteinase K at 55°C for 30 minutes. Beads were resuspended in TRIzol (Invitrogen 15596018), RNA extraction was performed immediately followed by RNA-sequencing.

Tandem Immunoprecipitation

After biotin washout, control and Actinin-BirA* iPSC-CMs were rinsed three times with room temperature PBS and then lysed in RIP buffer. Cleared lysates were incubated with Dynabeads Protein G (Invitrogen 10003D) pre-bound to either IgG antibody (MBL PM035) or IGF2BP2 antibody (MBL RN008P). Samples and beads were incubated on a rotator overnight at 4°C. The following day, the Protein G beads were washed three times with RIP buffer and then incubated for 12 hours with rotation at 4°C in RIP buffer containing IMP2 peptide (Genescript). The supernatant was then mixed with magnetic Dynabeads MyOne Streptavidin C1 beads (Invitrogen 65001), which were prewashed with PBS and RIP buffer. The following day, the streptavidin beads were three times washed with ice-cold RIP buffer – all steps were performed in a 4°C room. Samples were then incubated with Proteinase K at 55°C for 30 minutes. Beads were resuspended in TRIzol, RNA extraction was performed immediately followed by cDNA synthesis (outlined in Quantitative PCR section).

**Quantitative proteomics**

Bead digestion

After conducting streptavidin immunoprecipitation, streptavidin beads were snap frozen and sent to the Thermo Fisher Center for Multiplexed Proteomics (TCMP) at Harvard

Medical School. For each tandem mass tag (TMT) mass spectrometry experiment, three biological replicates per condition were submitted.

The beads were washed three times with 50 mM Tris buffer, pH 8.0 to remove any trace of detergents and unspecific binders followed by resuspension in 1M Urea, 50 mM Tris, pH 8.0 and initially digesting it with Trypsin (5 ng/μl) at 37°C for one hour under shaking. After initial trypsin incubation, samples were centrifuged briefly and the supernatant were collected in new tubes. Beads were further washed three times with 1M Urea, 50 mM Tris, pH 8.0 and washed fractions were pooled in the same tubes and left to digest overnight at room temperature. The following morning, digested peptides were reduced first with 5 mM TCEP, followed by alkylation with 10 mM Iodoacetamide, quenching alkylation with 5 mM DTT and finally quenching the digestion process with TFA. Acidified digested peptide were desalted over C18 stagetip following protocol described before[7]. Briefly, the tips were prepared placing a small disc of Empore material 3M in an ordinary pipette tip, preparing a single tip for each sample. Tips were cleaned and secured with Methanol, activated with 50% acetonitrile, 0.1% TFA, equilibrated with 0.1% TFA. Acidified digested sample was added to the column and finally washed twice with 0.1% TFA solution. Liquid was passed through the pipette tip with a centrifugation. Peptides were then eluted with 80% acetonitrile, 0.1% TFA buffer thrice and dried in a speedvac. Dried desalted nine peptides samples were reconstituted with 200 mM EPPS buffer, pH 8.0 and labelled with respective first 9 of a 10-plex tandem mass tag (TMT) reagent. The 9-plex labeling reactions were performed for 1 hour at room temperature. Modification of tyrosine residues with TMT was reversed by the addition of 5% hydroxyl amine for 15 minutes and the reaction was quenched with 0.5% TFA. Samples were combined, further desalted over stage-tip, finally eluted into an Autosampler Inserts (Thermo Scientific™), dried in a speedvac and reconstituted with 5% Acetonitrile-5% TFA for MS analysis.

Liquid chromatorgraphy-MS3 spectrometry (LC-MS/MS)

Labelled peptide sample from the previous step was analyzed with an LC-MS3 data collection strategy[8] on an Orbitrap Fusion mass spectrometer (Thermo Fisher Scientific) equipped with a Thermo Easy-nLC 1200 for online sample handling and peptide separations. Resuspended peptides from the previous step was loaded onto a 100 µm inner diameter fused-silica micro capillary with a needle tip pulled to an internal diameter less than 5 µm. The column was packed in-house to a length of 35 cm with a $C_{18}$ reverse phase resin (GP118 resin 1.8 µm, 120 Å, Sepax Technologies). The peptides were separated using a 180 min linear gradient from 5% to 42% buffer B (90% ACN + 0.1% formic acid) equilibrated with buffer A (5% ACN + 0.1% formic acid) at a flow rate of 500 nL/min across the column.

The scan sequence for the Fusion Orbitrap began with an MS1 spectrum (Orbitrap analysis, resolution 120,000, scan range of 350 - 1350 m/z, AGC target 1 x 106, maximum injection time 50 ms, dynamic exclusion of 90 seconds). The “Top10” precursors was selected for MS2 analysis, which consisted of CID (quadrupole isolation set at 0.7 Da and ion trap analysis, AGC $9 \times 10^3$, Collision Energy 35%, maximum injection time 80 ms). The top ten precursors from each MS2 scan were selected for MS3 analysis (synchronous precursor selection), in which precursors were fragmented by HCD prior to

Orbitrap analysis (Collision Energy 55%, max. AGC 1 × $10^5$, maximum injection time 120 ms, resolution 50,000 and isolation window set to 1.2 – 0.8).

LC-MS3 data analysis

A suite of in-house software tools were used for .RAW file processing and controlling peptide and protein level false discovery rates, assembling proteins from peptides, and protein quantification from peptides. MS/MS spectra were searched using the SEQUEST[9] algorithm against a Uniprot composite human database (human release 2017-10) with both the forward and reverse sequences. Database search criteria are as follows: tryptic with two missed cleavages, a precursor mass tolerance of 50 ppm, fragment ion mass tolerance of 1.0 Da, static alkylation of cysteine (57.02146 Da), static TMT labeling of lysine residues and N-termini of peptides (229.162932 Da), and variable oxidation of methionine (15.99491 Da). Peptide spectral matches were filtered to a 1% false discovery rate (FDR) using the target-decoy strategy combined with linear discriminant analysis. The proteins were filtered to a <1% FDR. TMT reporter ion intensities were measured using a 0.003 Da window around the theoretical m/z for each reporter ion in the MS3 scan. Proteins were quantified only from peptides with a summed SN threshold of >200 and MS2 isolation specificity of 0.5. Peptide spectral matches with poor quality MS3 spectra were excluded from quantitation (<200 summed signal-to-noise across 10 channels and <0.5 precursor isolation specificity).

Before analysis, proteins previously reported to non-specifically bind BirA* were removed[10]. For analysis, TMT raw intensity values underwent Log2-fold-change (L2FC) calculation and enrichment analysis. For actinin proteome analysis (Figure 1), data from experiments 1 and 2 were utilized to determine significantly enriched proteins by calculating the L2FC of proteins from Actinin-BirA* relative to Control. Proteins considered to be significantly enriched were those with L2FC ≥1 and false discovery rate (FDR) <0.05 (using two-way ANOVA followed by a two-stage linear step-up procedure of Benjamini, Krieger and Yekutieli to correct for multiple comparisons[11]). For sarcomere assembly actinin proteome analysis (Figure 2), data from experiment 2 was similarly analyzed to first determine enriched proteins using cTnT-WT-Actinin-BirA* relative to Control. Those enriched proteins were then analyzed for sarcomere-dependency status by calculating the L2FC of proteins from cTnT-WT-Actinin-BirA* relative to cTnT-KO-Actinin BirA*, with significant hits being defined as those with L2FC ≥1 and FDR <0.05. For visualization of data, Search Tool for the Retrieval of Interacting Genes/Proteins (STRING v. 11.0) was utilized for generation of interaction scores (proteins were not distinguished by isoform). Cytoscape (v.3.7.2) was utilized to generate interaction maps with an organic layout and the utilization of the CLUSTER and BINGO features for group clustering and appropriate GO term identification for each cluster. Quantitative proteomics data deposited to the ProteomeXchange Consortium identifier PXD018040.

**Quantitative PCR, RNA sequencing, and analysis**

cDNA was synthesized using Superscript III First-Strand synthesis (Invitrogen 18080-400). Gene-specific PCR primers were designed or identified from the literature and transcripts were quantified using Fast SYBR Green (Applied Biosystems 4385612) on a ViiA7 Real-Time PCR system (Applied Biosystems).

RNA Samples were sent for sequencing at the University of Connecticut Institute for Systems Genomics. RNA sequencing libraries were generated using Illumina TruSeq Stranded Total RNA library preparation (Ribo-Zero depletion for rRNA was utilized for

RIP-seq). Illumina NextSeq 500/550 sequencing was conducted with the v2.5 300 cycle reagent kit 9 (High Output). Estimated total single end reads per sample = 30-35M 150bp PE reads. Samples sequences were aligned with STAR to the hg38 human genome. In order to look at differential expression, DESeq2 (Bioconductor) was utilized. Gene Set Enrichment Analysis (GSEA) was utilized to determine GO terms for data sets. Top 10 GO terms are presented. Furthermore, for ETC complexes, glycolysis, and sarcomere gene sets – GSEA gene sets were utilized and those with an average ≥5 TPM value were included in the data set. TPM (transcripts per million) is a normalization technique for transcriptomic data to adjust for transcript count read depth and transcript length, such that all transcript counts are normalized by transcript length and then presented as a proportion out of 1 million counts. Genome sequencing datasets are deposited at GEO under accession numbers: GSE144805 and GSE144806.

**Protein Immunoblotting**

Samples of interest were lysed in RIPA buffer (Cell Signaling 9806) containing protease inhibitor cocktail (Roche 11836170001), 1 mM PMSF, and phosphatase inhibitor (Pierce A32957), unless noted otherwise. Protein lysate concentrations were normalized using Pierce BCA (Thermo 23225), and then reduced and denatured in sample buffer (Thermo 39000). Lysates were separated on Bio-Rad 4–20% Mini-PROTEAN TGX precast gels, transferred via Bio-Rad Trans-Blot Turbo onto PVDF membranes (Bio-Rad 1704272), blocked in TBS-T (TBS with 0.1% Tween-20) containing 5% BSA, and probed overnight at 4°C with primary antibody. The next day, blots were washed in TBS-T and probed with HRP-linked secondary antibody (Cell Signaling 7076; 7074; Streptavidin-HRP 3999S) for 1 hour. Signal detection was performed using ECL substrate (Thermo 34577; 34095) and a Bio-Rad ChemiDoc MP imaging system. Blot images were digitally processed and analyzed in ImageJ. The primary antibodies used were as follows: anti-actinin (Cell Signaling 6487), anti-OxPhos (Invitrogen 458199) anti-HA (Cell Signaling 14793), anti-IGF2BP2 (MBL RN008P), anti-PBCBP1 (Abcam 74793), anti-PCBP2 (MBL RN250P), anti-SERBP1 (Abcam 55993), anti-TCAP (BD 612328), and anti-MYOM (mMaC myomesin B4 was deposited to the DSHB by Perriard, J.-C.).

**Immunofluorescence and RNA FISH**

For standard immunofluorescence images, cells were fixed on coverslips (Fisherbrand 12-545-100) in PBS containing 4% paraformaldehyde (EMS 50-980-487) and then permeabilized and blocked in PBS-T (PBS containing 0.1% Triton-X) with 1% BSA (Fisher BP1605100). Cells were probed with primary antibody in PBS-T with BSA overnight at 4°C. The following primary antibodies were used:, anti-Actinin (Sigma A7811) and anti-IGF2BP2 (Bethyl A303-316A). The next day coverslips were washed in PBS-T, incubated for 1 hour at room temperature with 1 µg/mL DAPI (Invitrogen D1306) and appropriate secondary antibody (Invitrogen A-11005 or A-11008) or streptavidin-488 (Invitrogen S11223), washed, and mounted onto slides (Corning 2948) in ProLong Diamond mountant (Invitrogen P36965). An Andor Dragonfly 500 confocal microscope system using a Zyla sCMOS camera and a Leica DMi8 63x oil immersion lens was used for imaging slides. Andor Fusion software was used to acquire images, which were converted to OME-TIFF format using Bitplane Imaris and analyzed in ImageJ.

ViewRNA direct fluorescence RNA in situ hybridization was conducted using the manufacturer's protocol (Invitrogen 19887). The following human probes were utilized: *NDUFA1* (Invitrogen VA6-3172657), *NDUFA8* (Invitrogen VA1-3003564), and *TTN*

(Invitrogen VA6-3173894). Final step of coverslip mounting and image acquisition are described above. For distance quantification, Imaris software was utilized. Images first underwent background subtraction to minimize background signal and allow for clear puncta visualization. The distance transformation tool on Imaris was used to quantify distance (in microns) of RNA puncta and actinin structures.

**Mammalian-Two Hybrid**

Actinin (full-length and domains), IGF2BP2 (full-length and domains), PCBP1, PBCBP2, and SERBP1 were amplified from human cDNA and cloned into the appropriate vectors (pACT or pBIND) provided by the CheckMate Mammalian Two-Hybrid kit (Promega E2440). The appropriate combination of edited pACT and pBIND vectors, in addition to the kit-provided pGluc5 vector, were transfected using PEI into HEK 293T cells (ATCC CRL-3216), as outlined in the manufacturer's protocol. In addition, the kit-provided positive and negative controls were transfected simultaneously for all experiments to verify proper signal production. Luciferase and Renilla activity were measured using manufacturer's protocol (Promega E1910) utilizing BioTek's Synergy 2 multi-mode microplate reader.

**Lentivirus Production**

HEK 293T cells (ATCC CRL-3216) were seeded onto 150 mm plates in 20 mL DMEM (Gibco 11965092) supplemented with 10% FBS (Gemini 100-106), GlutaMAX (Gibco 35050061), and 1 mM sodium pyruvate (Gibco 11360070). HEK 293T cells were transfected with 18ug of lentiviral transfer vector, 12 µg psPAX2 (Addgene 12260), and 6 µg pCMV-VSV-G (Addgene 8454). Media was replenished the following day and virus-containing media was harvested at 48, 72, and 96 hours post-transfection, followed by concentration using PEG-6000 as previously described[12]. For titer determination - iPSCs were transduced with a serial dilution of lentivirus, followed by treating with appropriate antibiotic (1µg/ml puromycin), and counting resistant colony-forming units.

**Polysome profiling**

IPSC-CMs at day 25 were treated with 0.1 mg/ml cycloheximide at 37 °C for 10 minutes. Cells were washed with 0.1 mg/ml cycloheximide in PBS and lysed in polysome lysis buffer (10 mM HEPES ph7.3, 150 mM KCl, 10 mM $MgCl_2$, 1 mM DTT, 100ug/ml cycloheximide, 2% NP-40, protease inhibitor mini tablet, 6U/ml RNase inhibitor). Samples were kept on ice for 5 minutes during lysis and then centrifuged at 13000g for 10 minutes at 4 °C to remove debris. The BioComp Gradient Station ip Model 153 was used to prepare sucrose gradients. 15% and 50% sucrose solutions (sucrose diluted in 10 mM HEPES pH 7.3, 150 mM KCl, 10 mM $MgCl_2$, 100 ug/ml cycloheximide, 1mM DTT) were utilized. Supernatants from lysed cells were added to sucrose gradients and centrifuged for 150 minutes at 4°C at 120000g (SW41 rotor, Beckman Coulter Optima XPN-100 Ultracentrifuge). BioComp Gradient Station and TRIAX software were utilized for separation of sucrose gradient fractions. Fractions were collected, resuspended in TRIzol and RNA extraction was conducted followed by RNA-Sequencing

**Individual Gene Knockdown**

Lentiviral-based hairpins were used to individually knockdown genes directly in iPSC-CMs. iPSC-CMs were transduced with lentivirus expressing hairpin at a MOI of 2 in RPMI-B27. Media was replenished the following day and cells were analyzed 7-10 days post transduction.

**Flow Cytometry Analysis**

BD Biosciences FACSymphony A5 was utilized to perform all flow cytometry experiments. Cells were stained in suspension for 30 minutes at 37°C with 10 μM Hoechst 33342 (Thermo 62249), 200 nM MitoTracker Green (Invitrogen M7514), and 500 nM TO-PRO-3 (Invitrogen T3605). Cells were gated based on forward-scatter and side-scatter, followed by gating of TO-PRO-3 to identify live cells. Hoechst was used to gate for single cells, followed by subsequent analysis of mitochondrial content (MitoTracker). Final analysis was performed using FlowJo software.

**Mito Stress Test**

30,000 wildtype CMs were plated on Seahorse XFe96 Cell Culture Microplates (Agilent 101085-004) and lentiviral shRNAs (Scramble and IGF2BP2) were added the next day. The cells were cultured for 7 days before performing the Seahorse XF Cell Mito Stress Test (Agilent 103015-100) according to manufacturer's protocol. The following final concentrations of compounds were injected: oligomycin (1 μM), FCCP (1 μM), and rotenone/antimycin A (0.5 μM).

**Cardiac microtissue assay**

Cardiac tissues were generated as previously described[2, 6]. Briefly, polydimethylsiloxane (PDMS) (Corning Sylgard 184) cantilever devices were molded from SU-8 masters, which were embedded with fluorescent microbeads (Thermo F8820). Singularized iPSC-CMs were mixed with human cardiac fibroblasts and spun into PDMS devices containing a collagen-based ECM. For knockdown experiments, lentiviral shRNAs were added to iPSC-CMs 7 days prior to generation of tissues. Tissue function was measured once tissues compacted and were visibly pulling the cantilevers. For acquisition of functional data, tissues were stimulated at 1 Hz with a C-Pace EP stimulator (IonOptix) with platinum wire electrodes and fluorescence images were taken at 25 Hz with a Andor Dragonfly microscope equipped with enclosed live-cell chamber (Okolabs). Displacement of fluorescent microbeads was tracked using the ImageJ ParticleTracker plug-in and twitch force was calculated. The cantilever spring constants were determined using the elastic modulus of PDMS and the dimensions of the tissue gauge devices previously described[13].

**Statistical Analysis**

Obtained data were analyzed in Microsoft Excel and graphed using either GraphPad Prism or R. Data are presented as mean ± standard error of the mean (SEM). Statistical comparisons were conducted via Student's t-test or one-way ANOVA followed by multiple comparisons using Holm-Sidak correction, where appropriate. Statistical significance was defined by $p \leq 0.05$ (*), $p \leq 0.01$ (**), $p \leq 0.001$ (***), and $p \leq 0.0001$ (****).

#### Supplemental Figure Legends

**Figure 1. BioID to establish actinin proximity partners in iPSC-CMs. A)** Overview of gene-targeting strategy to generate an in-frame knock-in of BirA*-HA at the *ACTN2* locus in iPSCs. CRISPR/Cas9 and homology-directed repair from a donor vector containing BirA*-HA and 800bp *ACTN2* homology arms was utilized. **B)** Actinin-BirA* cardiac microtissues have similar twitch force compared to controls. Data are n≥10 microtissues. (scale bar-150μm). **C)** Representative protein blot probed with streptavidin-HRP to optimize biotin labeling after 24 hours of biotin supplementation up to 50 μM. **D)** Confocal micrograph of Control (non-BirA*) iPSC-CMs decorated with antibodies to actinin (red), streptavidin-AF488 (green) and DAPI DNA co-stain (blue) showing diffuse streptavidin staining in the absence of Actinin-BirA* (scale bars, main image-10μm). **E)** Complete STRING network map of Actinin proximity proteins. Data are mean ± SEM; significance assessed by Student's t-test (b) and defined by $P > 0.05$ (ns).

**Figure 2. BioID through sarcomere assembly. A)** Representative immunoblot of cTnT-KO and cTnT-WT iPSC-CMs lysates probed for cardiac troponin T (cTnT), actinin and GAPDH control. **B)** Representative protein blot of cTnT-KO and cTnT-WT Actinin-BirA* lysates probed with streptavidin-HRP**.**

**Figure 3. RIP-seq to identify RNA transcripts. A)** Heatmap displaying relative expression of ETC Complexes (I-V) genes in Control and Actinin-BirA* RIP-seq samples. **B)** Heatmap displaying relative expression of glycolysis genes in Control vs. Actinin-BirA* RIP-seq samples. **C)** Heatmap displaying relative expression of sarcomere thin and thick filament genes in Control vs. Actinin-BirA* RIP-seq samples. **D)** qPCR validation of candidate RIP-seq hits. Data are n=4. **E**) Representative image from Imaris utilizing distance transformation tool to determine distance between actinin puncta (red) and mRNA puncta (multi-colored). **F**) Representative confocal micrograph of iPSC-CMs decorated with antibodies to actinin (red), DAPI DNA co-stain (blue), and RNA FISH probe against *TTN* (purple). (scale bars, main image-10μm, inset-5μm). **G)** Representative confocal micrograph of iPSC-CMs decorated with antibodies to actinin (red), DAPI DNA co-stain (blue), and RNA FISH probe against *NDUFA8* (cyan). (scale bars, main image-10μm, inset-5μm). **H)** RNA FISH of ETC Complex I components NDUFA1 and NDUFA8 that are both in proximity to actinin but not overlapped (scale bars, main image-10μm, inset-5μm). Data are mean ± SEM; significance assessed by two-way ANOVA using Holm-Sidak correction for multiple comparisons (d) and defined by $P < 0.05$ (*), $P \leq 0.01$ (**), $P \leq 0.0001$ (****).

**Figure 4. RNA-binding proteins interact with actinin. A)** Representative immunoblot probed with antibodies to PCPB1, PCPB2 and SERBP1 in streptavidin affinity-purified lysates. **B)** Representative immunoblot probed with antibodies to PCPB1, PCPB2 and SERBP1 in anti-HA immunoprecipitated lysates. **C)** M2H (conducted in HEK 293T cells) results demonstrating that IGF2BP2 but not PCBP1, PCPB2 and SERBP1 interact with full-length actinin. **D)** Summary overview of RBP and actinin interaction data. Data are n=3; mean ± SEM; significance assessed by one-way ANOVA using Holm-Sidak correction for multiple comparisons (c) and defined by $P \leq 0.0001$ (****).

**Figure 5. IGF2BP2 localizes ETC transcripts to actinin. A)** Overview of tandem-IP process (left) and qPCR results of IgG vs. IGF2BP2 IP (right) for candidate ETC transcripts. Data are n=3; mean ± SEM; significance assessed by Student's t-test (a) and defined by $P < 0.05$ (*). **B)** Example of gating strategy utilized for analysis of mitochondrial content for Main Figure 5e.

**Table 1**. Raw TMT intensity data for Experiment 1.

**Table 2**. Raw TMT intensity data for Experiment 2.

**Table 3.** Data for actinin-proximity proteins.

**Table 4.** Data for sarcomere assembly proteins.

**Table 5.** RIP-seq data and GO Terms.

**Table 6.** Overlay of RIP-seq and Polysome-seq data and GO terms.

**Table 7.** CRISPR and primer sequences.

**Movie 1.** Control Actinin-BirA* iPSC-CMs paced at 1Hz. Magnification-5X. Scale bar-150μm.

#### Supplemental Figure 1

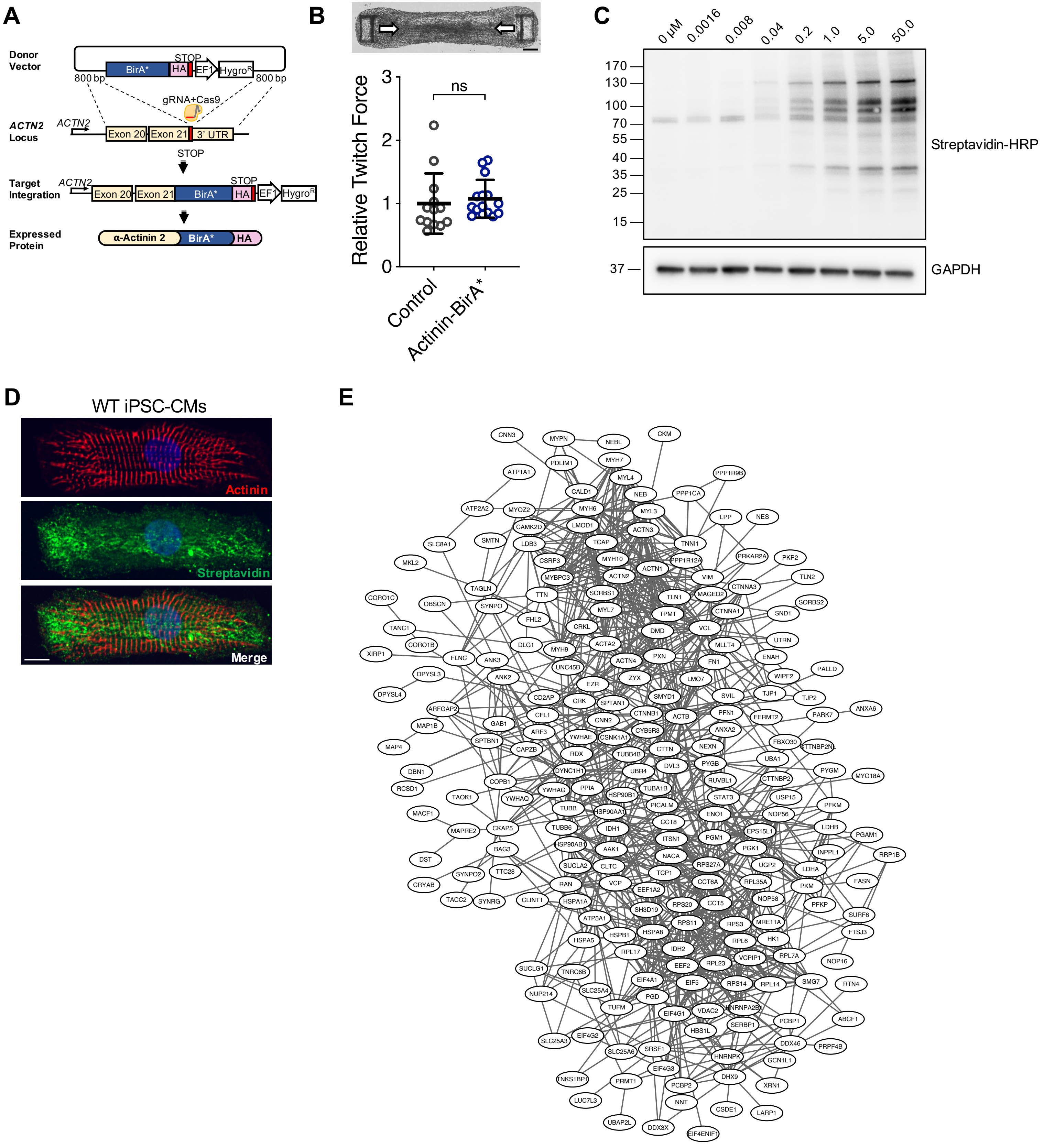

#### Supplemental Figure 2

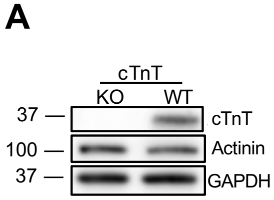


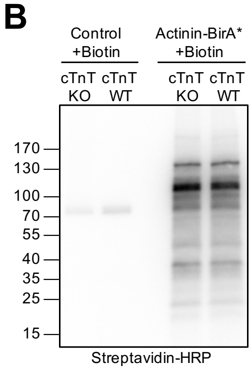

#### Supplemental Figure 3

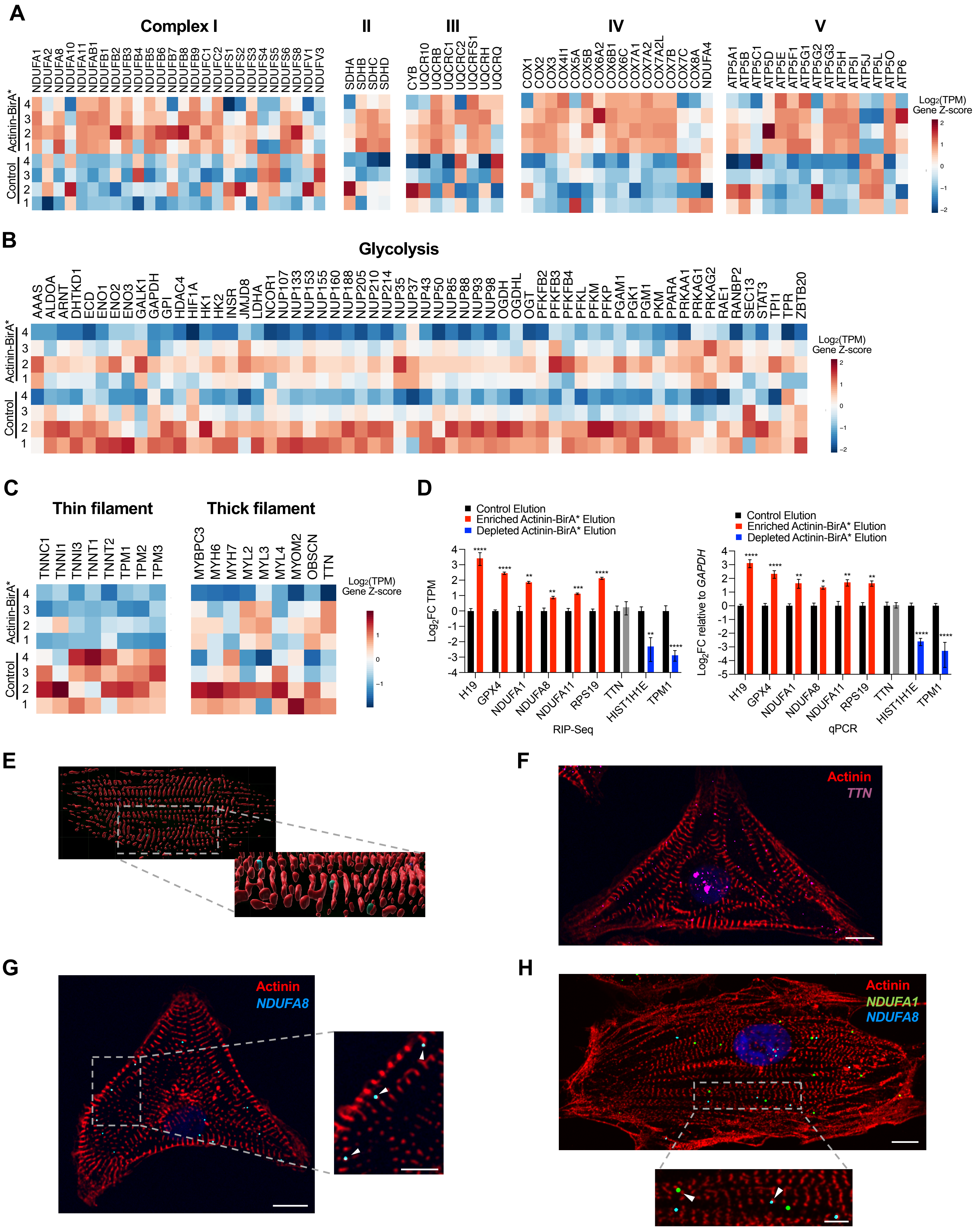

#### Supplemental Figure 4

**A**

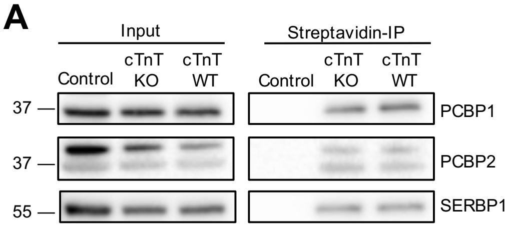


**B**

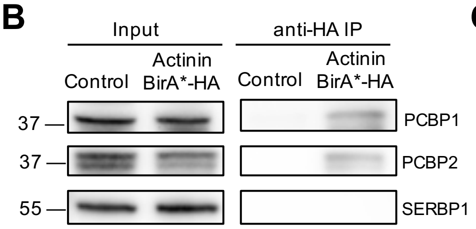


**C**

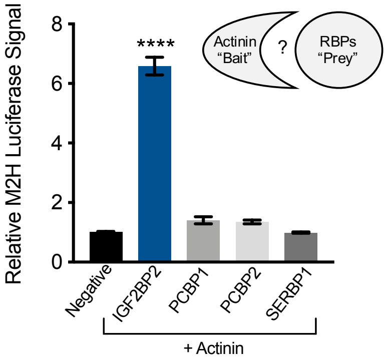


**D**

| **RNA Binding Protein** | **Method** | | | |
|---|---|---|---|---|
| | TMT | SP-IP | HA-IP | M2H |
| IGF2BP2 | + | + | + | + |
| PCBP1 | + | + | + | - |
| PCBP2 | + | + | + | - |
| SERBP1 | + | + | - | - |

### Supplemental Figure 5

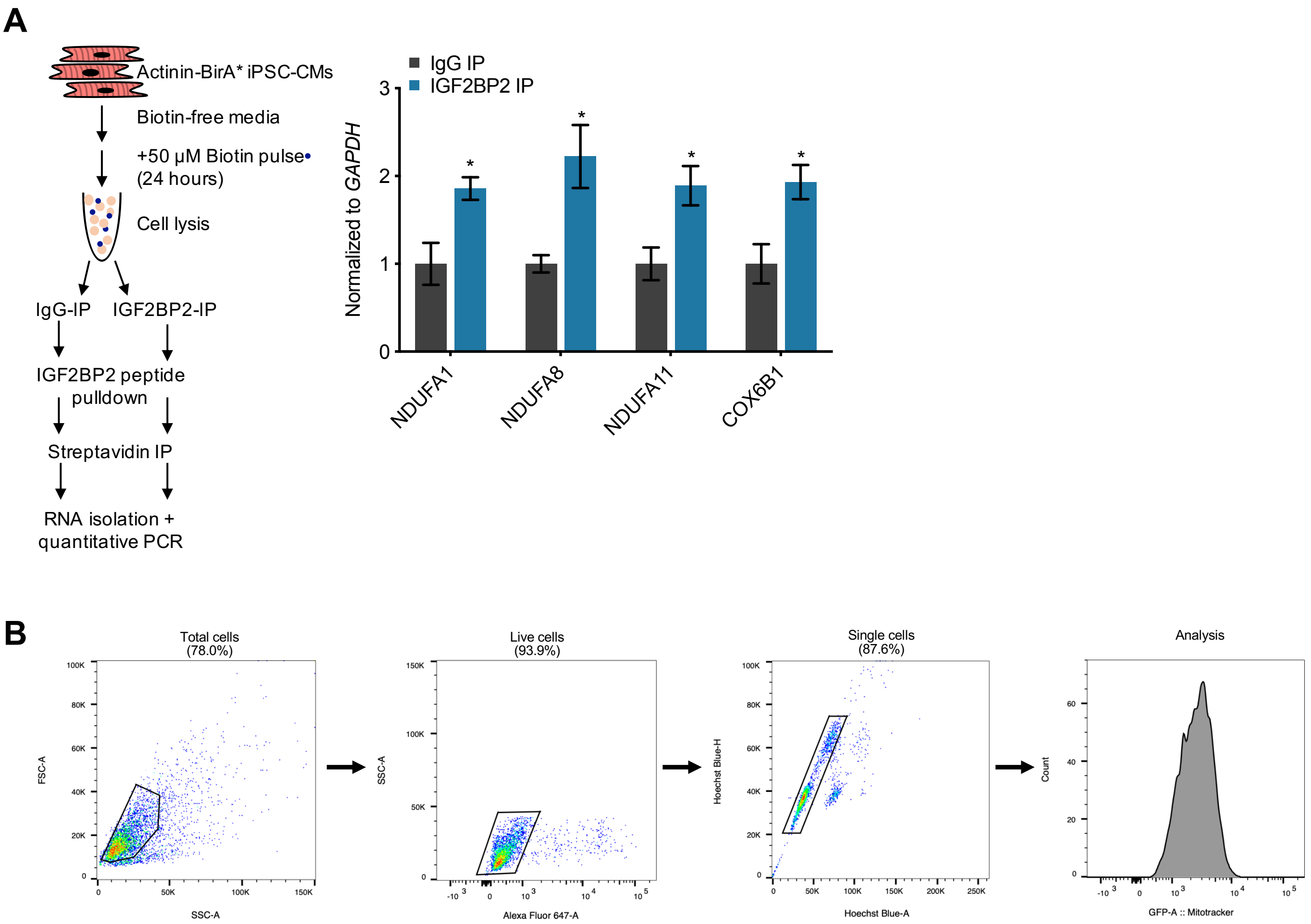
